# Multi-Site Reproducibility Study of 3D High-Content Analysis with Dual-View Oblique Plane Microscopy

**DOI:** 10.64898/2026.06.29.735376

**Authors:** Yuriy Alexandrov, Mar Arias-Garcia, Chris Bakal, Eduard Batlle, Vicky Bousgouni, Neil Carragher, Julien Colombelli, Jayne Culley, Nathan Curry, Lucas Dent, Chris Dunsby, Liuba Dvinskikh, Edwin Garcia, Nikolaos N. Giakoumakis, Nils Gustafsson, Montserrat Llanses, Martin Lee, Kanad N. Mandke, Dan Marks, Iain McNeish, Colin D. H. Ratcliffe, Erik Sahai, Hugh Sparks, Theresa Suckert

**Affiliations:** Light Community, Department of Physics, Imperial College London, South Kensington Campus, London, SW7 2AZ, United Kingdom; The Francis Crick Institute, 1 Midland Road, London, NW1 1AT, United Kingdom; Institute of Cancer Research (ICR), 123 Old Brompton Road, London, SW7 3RP, United Kingdom; Institute for Research in Biomedicine (IRB Barcelona), Carrer de Baldiri Reixac, 10, 08028 Barcelona, Spain; Cancer Research UK Scotland Centre, Institute of Genetics and Cancer, University of Edinburgh, Edinburgh, EH4 2XU, United Kingdom; Ovarian Cancer Action Research Centre, Department of Surgery and Cancer, Imperial College London, London W12 0NN, UK

## Abstract

High content imaging is being applied to achieve quantitative fluorescence readouts in increasingly complex 3-dimensional (3D) cell culture models such as spheroids and organoids. Compared to conventional 2D assays, 3D assays better represent biological heterogeneity but require more complex sample preparation, 3D imaging and 3D image analysis that can affect the accuracy and precision of such assays. We used spheroids formed from the NRAS-activated melanoma cell line 19161 modified to express an ERK kinase translocation reporter (KTR) as an exemplar 3D phenotypic assay carried out in 96-well plates. The spheroids were treated with the ERK activator TPA and a range of concentrations of the MEK inhibitor Binimetinib. 3D live-cell imaging with sub-cellular spatial resolution was performed using a dual-view oblique plane microscope (dOPM) – a form of single-objective light-sheet microscope – and the experiment was performed separately at 4 different institutes. The results were analysed using an identical 3D analysis pipeline and parameters. We assessed the variation in assay readout using a linear mixed effects model. Random variance at the well level was negligible (SD = 0.0048 relative to range of KTR biosensor readout at reference site of 0.17), indicating low technical noise. Treatment effects were dose-dependent and highly statistically significant compared to DMSO control across all sites (Dunnett-corrected *p* < 0.001). The range in KTR readout between the minimum (3.5 μM Binimetinib) and maximum (100 nM TPA) treatments varied between 59 to 96% relative to the reference site. Measured bias in KTR readout between sites was between 6 and 12% of the range of the reference site. This study quantifies the reproducibility of a 3D live spheroid-based assay employing a fluorescence biosensor requiring readout out at the per-cell level using the dOPM platform and discusses areas where experimental protocol could be improved in the future to further improve reproducibility.

## Introduction

High content imaging (HCI) of arrays of cells grown in 2D in a multi-well plate is a standard tool in the biomedical sciences for studying the response of cells to a range of different experimental conditions including high throughput screening of multiple genetic or pharmacological perturbations [1, 2]. The use of 2D cell culture is convenient and facilitates optical imaging readouts at scale using either wide-field epi-fluorescence microscopy or an optically sectioned imaging method such as confocal or spinning-disk confocal microscopy to reject out-of-focus signal, increase the signal-to-background ratio and to permit more accurate quantification of cellular and sub-cellular parameters [3].

There is increasing interest to use more biologically relevant 3D cell cultures such as spheroids, organoids and organ-on-chip technologies in HCI [4–8]. In cancer biology, this shift is driven by the need to better model tumour complexity, heterogeneity and localised signalling environments [9] and reduce high attrition rates of new candidate drugs in clinical trials. However, these 3D structures introduce additional experimental considerations, including how to form the spheroids/organoids reliably, how best to image these more optically challenging samples [10] and how best to analyse the resulting data.

For the highest resolution images, it is necessary to prepare spheroids in plates with optical quality coverslip bases. For fixed endpoint measurements, spheroids can be grown using conventional methods and then fixed and transferred into the imaging plate. For live cell imaging, spheroids can be grown in low adherence flat-bottomed plates, cells can be seeded into gels that either fill or form domes in the bottom of each well [11], or can be printed in arrays of gel spots [12]. Centrifugation is sometimes used prior to gelation to force cells nearer to the coverslip. Cells may also be grown in 3D gel cavities printed onto an optically flat base, e.g. [13].

HCI of spheroids or organoids can be carried out using automated wide-field microscopy or confocal/spinning disk confocal microscopy at one or a few z-positions. If multiple z-positions are acquired, then a z-projection of the data may be taken so that the analysis and quantification can be performed in 2D. These approaches yield bulk readouts of spheroid/organoid response but may not provide sufficient resolution to read out cell-wise or sub-cellular features [14].

Automated 3D imaging is required to read out more complex parameters and necessitates the use of a fully 3D image analysis pipeline. The imaging can be achieved using e.g. confocal microscopy, spinning-disk confocal microscopy or light-sheet fluorescence microscopy (LSFM). Conventional confocal microscopy can be too slow when needing to image 10s or 100s of fields-of-view in 3D within a reasonable timescale, often making spinning-disk confocal [3], line-scanning confocal [15] or LSFM more appropriate. LSFM of arrays of 10s of samples can be achieved using dual-objective LSFM systems where the samples are arrayed in a capillary tube [16], v-shaped transparent polymer foil troughs [17] or conical mounts [18]. Larger arrays of samples can be imaged using open-top light sheet microscopy [19, 20] or diSPIM [12], although approaches where samples share a common reservoir of culture medium are not compatible with testing a range of distinct treatment conditions in the same experiment [12, 20]. HCI LSFM can also be achieved using modified multiwell plates incorporating a mirror for light-sheet illumination [21] or by adding the mirror via an atomic-force microscope cantilever [22]. Another approach is to use oblique plane microscopy (OPM), which is a single-objective LSFM method that provides light-sheet illumination and fluorescence collection through the same primary objective lens and can be implemented on a conventional fluorescence microscope frame [23–25]. Dual-view OPM (dOPM) is a variant of OPM that enables two image volumes to be acquired with different light sheet illumination angles; the two views can subsequently be fused in software to reduce light-sheet shadowing artefacts and to provide a more isotropic spatial resolution [26, 27]. Light-field microscopy can provide volumetric images many times per second [28], however it works best with sparsely labelled samples and spatial resolution is traded off against field of view [10]. Two-photon microscopy provides the ability to image deeper into such samples, but the deepest imaging is obtained with single-point scanning which has image acquisition times similar to confocal microscopy.

As assays with full 3D imaging and 3D analysis become more widely used there is a need to understand how reproducible such assays, and the HCI platforms used, can be. When developing such assays, parameter choices in spheroid preparation [29] and image analysis [2] impact the reproducibility of the assay. A gold standard for understanding reproducibility in laboratory and clinical research are multi-site studies, which allow intra- and inter-site variability to be assessed [30, 31].

In this paper, we developed an exemplar 3D HCI assay read out at the single cell level in response to acute drug treatment using dOPM. Measurements were performed across four laboratory sites: The Francis Crick Institute in London, the Institute of Genetics and Cancer at the University of Edinburgh, The Institute of Cancer Research in London and the IRB Barcelona. We used a cancer spheroid assay in conventional 96-well plates seeding cells directly into basement-membrane extract gel. Spheroids were formed using an NRAS-activated mouse melanoma cell line engineered to express an ERK kinase translocation reporter (KTR). An exemplar dose-response assay was developed using a clinical MEK inhibitor (Binimetinib) and an upstream activator of MEK (TPA). We present the experimental protocol, biological effects, and quantify the variability in assay results both within and between sites.

### Protocol development and overview

We set out to develop a 3D spheroid-based assay that could be used in a test of assay reproducibility across 4 sites and be compatible with the working distance (280 μm) and field of view (∼200 μm) of our dOPM system [27]. This section provides an overview of the protocol and its development; full details are given in the Methods section.

We chose to base the assay on a pre-existing mouse NRAS activated melanocyte cell line (19161) [32, 33] modified at the ICR to express an ERK-KTR-mRuby2, EGFP-CAAX and H2B-iRFP670. The ERK-KTR [34] is used as the readout in the assay because NRAS driven melanoma generates constitutive activation of ERK1/2 via MEK1/2 [35]. This provides a matched readout of MEK pathway activity following treatment with a clinical MEK inhibitor (Binimetinib) in live single cells within a melanoma spheroid. The H2B-iRFP670 is used to count cells and to generate the nuclear segmentation mask required to quantify the biosensor readout.

The cell line was expanded at the ICR and then shipped to each of the other 3 partner sites. All sites purchased separate reagents from the same manufacturers using identical product codes. Therefore, the assay includes the effects of reagent batch-to-batch variability.

The assay was prepared by plating a suspension of single cells in 95% basement membrane extract (BME) into wells of a 96-well plate, see Figure 1. Once set, the BME gel was covered with 2% BME in culture medium. The spheroids were then grown over a period of 8 days. This growth method was chosen as a convenient and low-cost approach to generate spheroids within the working distance of the 60x water immersion microscope objective used for imaging.

**Figure 1:**
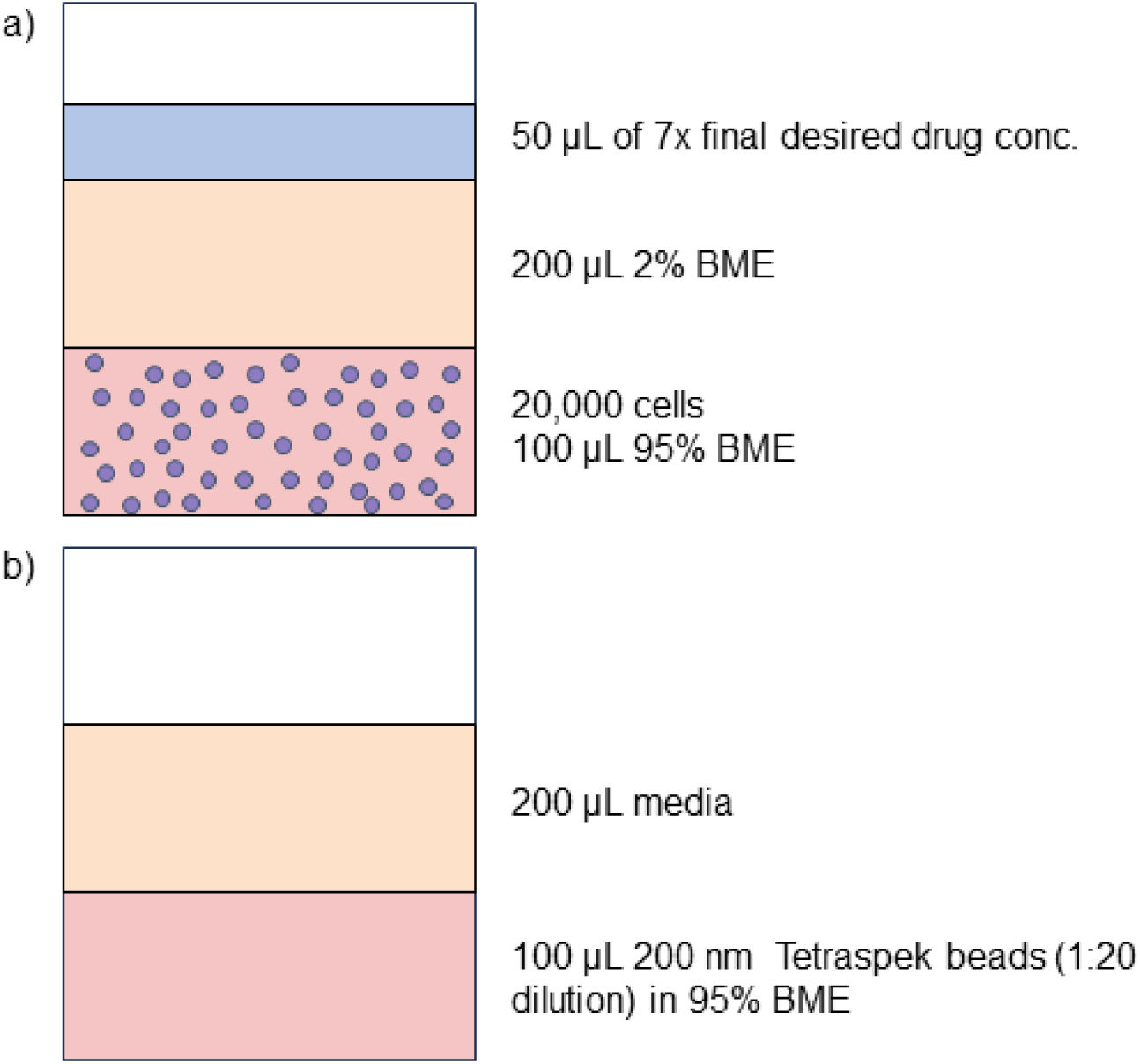
(a) Mouse NRAS-expressing melanoma spheroids in 95% Cultrex/BME with 2% BME media overlay and drug-containing media added on top. (b) Bead well includes 200 nm TetraSpeck beads in 95% BME for dual-view registration in dOPM imaging.

We chose to use 4 concentrations of the MEK inhibitor Binimetinib [36–38] plus vehicle (DMSO)-only treated spheroids in the assay. We also included an ERK activating treatment (TPA) as a further control and to establish the dynamic range of the ERK reporter. TPA is known to activate ERK via protein kinase C and MEK [39, 40]. The plate map for the assay is shown in Figure 2(a). Drugs were added 7 hours prior to start of imaging.

**Figure 2:**
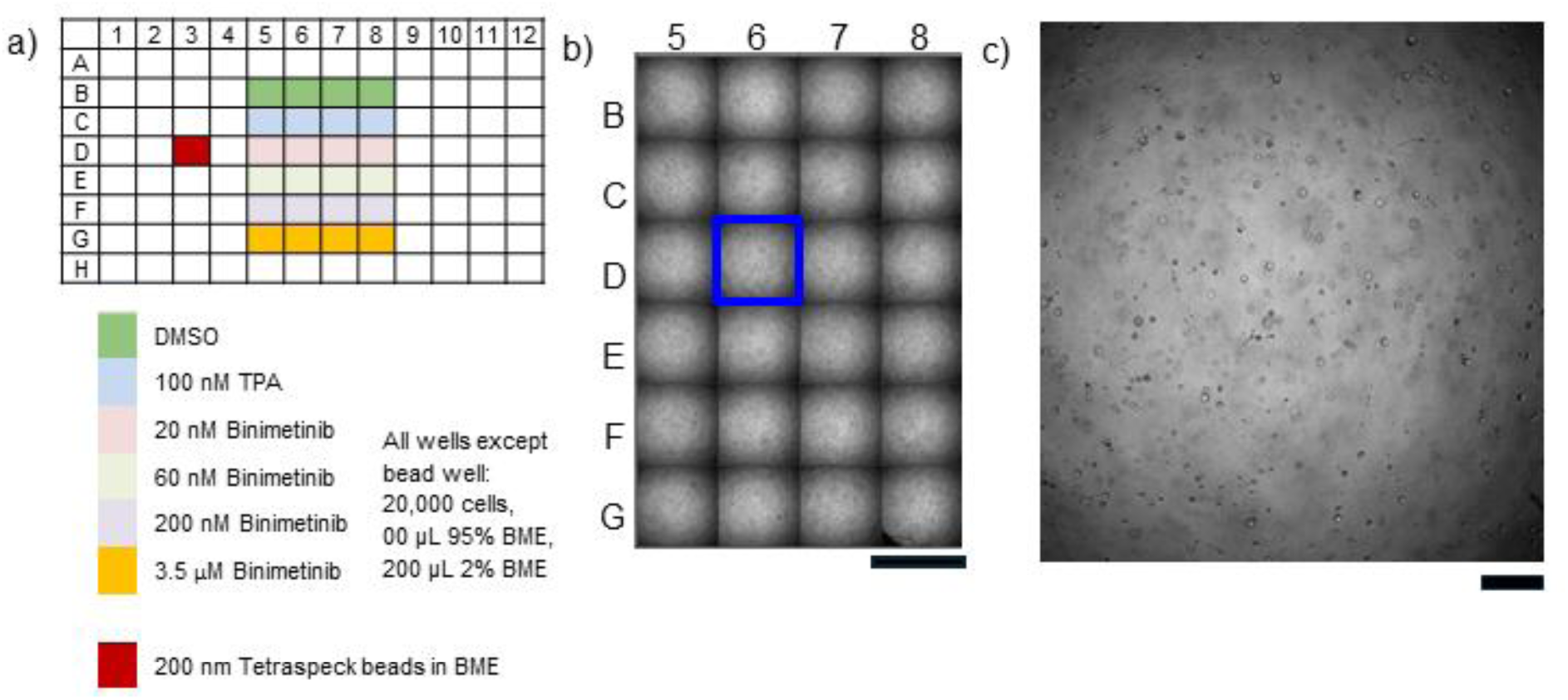
a) plate map. All spheroid-containing wells seeded with 20,000 cells, 100 μl 95 BME and overlaid with 200 μl 2% BME. b) 4× brightfield overview imaging of all spheroid-containing wells from a representative plate, scale bar 5 mm. c) Larger version of 4x brightfield overview of well D6, scale bar 500 μm.

As the initial cells are dispersed throughout the gel when seeded, spheroids are formed randomly through the gel. We therefore employed an automatic pre-find procedure that attempts to locate 10 spheroids within the dOPM microscope objective working distance in each well.

For data analysis, we chose to use a classical nuclear segmentation routine employing two-scale nonlinear top-hat filtering, binarisation and a watershed algorithm to separate touching nuclei. The nuclear regions of interest (ROIs) were then expanded to generate a 3D shell in the cytosol around each nucleus to sample the cytoplasmic biosensor signal. The KTR readout for each cell was then calculated as a ratio of the mean cytoplasmic intensity to the sum of the mean intensities of the cytoplasmic and nuclear regions.

## Results

The assay protocol developed was run independently at the four sites. The raw data generated is openly available: https://www.ebi.ac.uk/biostudies/bioimages/studies/S-BIAD3591.

Exemplar epifluorescence imaging during spheroid H2B-iRFP670 epifluorescence pre-find using a 20× air immersion objective is shown in Figure 3. Panel (b) shows the results of the Python segmentation with spheroids selected for imaging shown in yellow. Panel (c) shows epifluorescence imaging of the 10 centred in-focus spheroids automatically selected for imaging.

**Figure 3:**
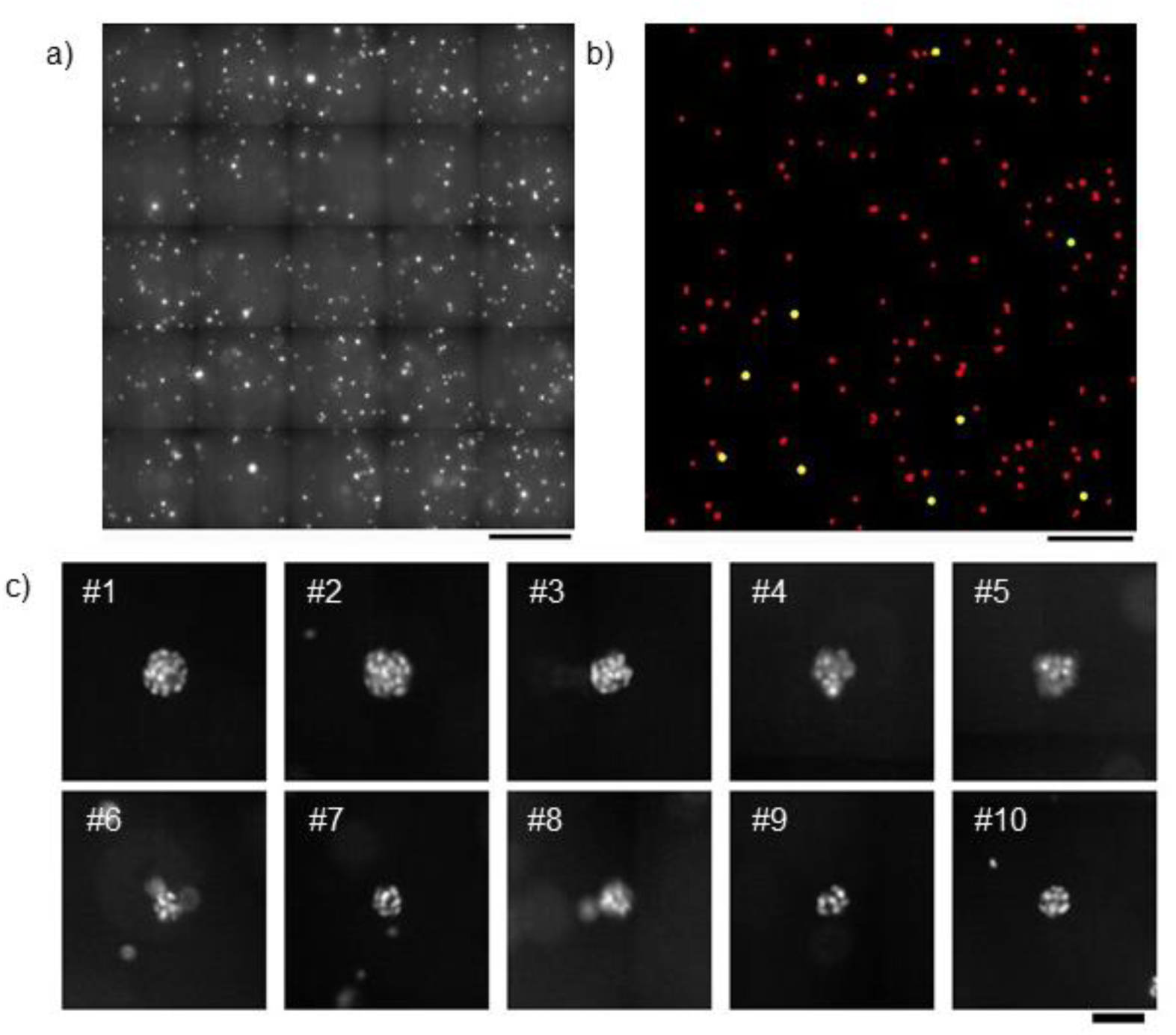
Wide-field epi-fluorescence prefind of spheroids with 20× objective showing exemplar well F6 treated with 20 nM Binimetinib (Step 3 of Table 7). (a) Epifluorescence tile scan of the H2B channel used for spheroid detection. Maximum intensity projection (MIP) of a 7-image stack acquired with 25 µm z-step. (b) Generated segmentation mask, where detected spheroid ROIs are marked in red and ROIs selected for dOPM imaging are marked in yellow. (c) Cropped images from the same 20x data shown in a) of the 10 selected spheroids at their estimated plane of best focus Scale bars: (a, b) 500 µm, (c) 50 µm.

Examples of the dOPM imaging are shown in Figure 4 for 3.5 µM Binimetinib (panel (b)) and 100 nM TPA (panel (c)). The ERK-KTR fluorescence signal is predominantly localised in the nucleus in (b) and in the cytoplasm in (c) as expected. 3D segmentation of the nuclei using the H2B-iRFP670 channel is shown in panel (d) together with the nuclear shell regions of interest in (e).

**Figure 4:**
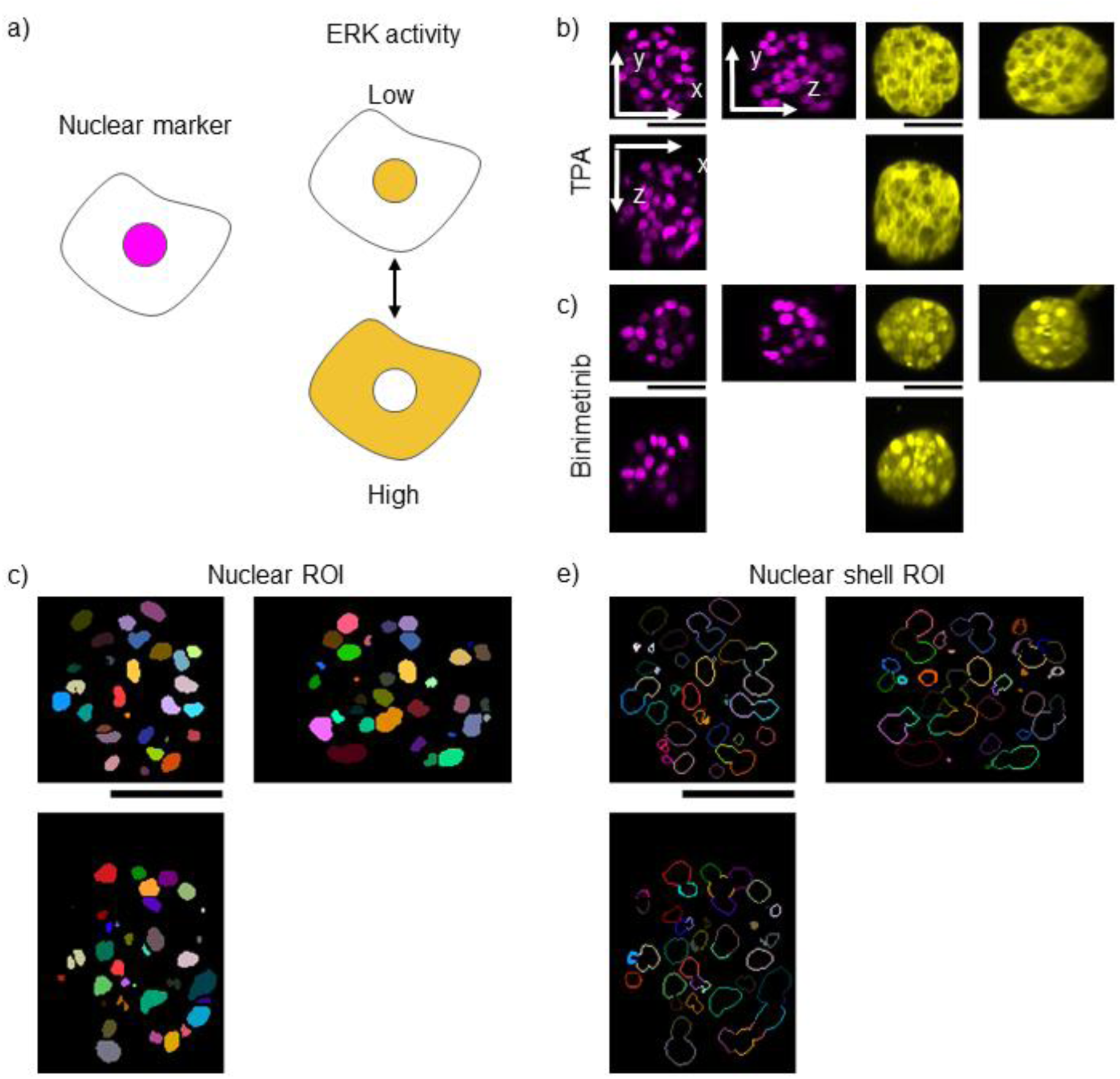
Single-cell ERK activity measured in 3D in melanoma spheroids (Step 5 of Table 7). (a) Cartoon of cellular localisation of nuclear marker H2B-iRFP670 (magenta) and ERK-KTR-mRuby2 biosensor (yellow). (b-e) Orthogonal slices (XY, ZY, XZ) through the central planes of exemplar dOPM image volumes of melanoma spheroids. (b) Well C7, 3.5 µM Binimetinib-treated spheroid with low ERK activity with nuclear-localized ERK-KTR-mRuby2. (c) Well G7, 100 nM TPA-treated spheroid with high ERK activity, with cytoplasmic localization of the biosensor. In (b&c) the H2B-iRFP670 signal is shown in magenta on the left and the ERK-KTR-mRuby2 signal is shown in yellow on the right. (d) nuclear ROIs generated from volume shown in (c). (e) Corresponding nuclear shell ROIs from volume shown in (c). Scale bars 50 µm.

The results of the assay replicated across 4 sites are shown in Figure 5, with montages of individual spheroid data shown in Supplementary Figure 1. The spatial distribution of the data is illustrated in Supplementary Figure 2. Figure 5(a) shows the median response of the KTR readout for each site calculated at the spheroid level. A similar trend is seen for all sites, and the differences are explored in more detail below. Figure 5(b) shows the median number of cells per spheroid calculated at the spheroid level. The size of the spheroids is reasonably consistent between conditions – i.e. across the well – as expected given the drug treatment time of 7 hours. The spheroids in site 1 had approximately double the number of detected cells relative to the other 3 sites. The raw signal levels for the nucleus (H2B-iRFP670), ERK-KTR-mRuby2 nucleus and ERK-KTR-mRuby2 cytoplasmic collar ROIs are shown in Supplementary Figure 3. Sites 3 and 4 had reasonably consistent nuclear signal, with the signal from site 2 being ∼3-fold lower and the signal from site 1 being ∼2-fold higher. For the nuclear and cytoplasmic collar ERK-KTR-mRuby2 signals, the signal levels from sites 2 and 4 were reasonably consistent with sites 3 being lower and site 4 being higher. The signals from sites 2 and 4 for the 100 nM TPA condition were noticeably lower compared to sites 1 and 3, but this did not noticeably affect the ratiometric KTR readout compared to the other sites.

**Figure 5:**
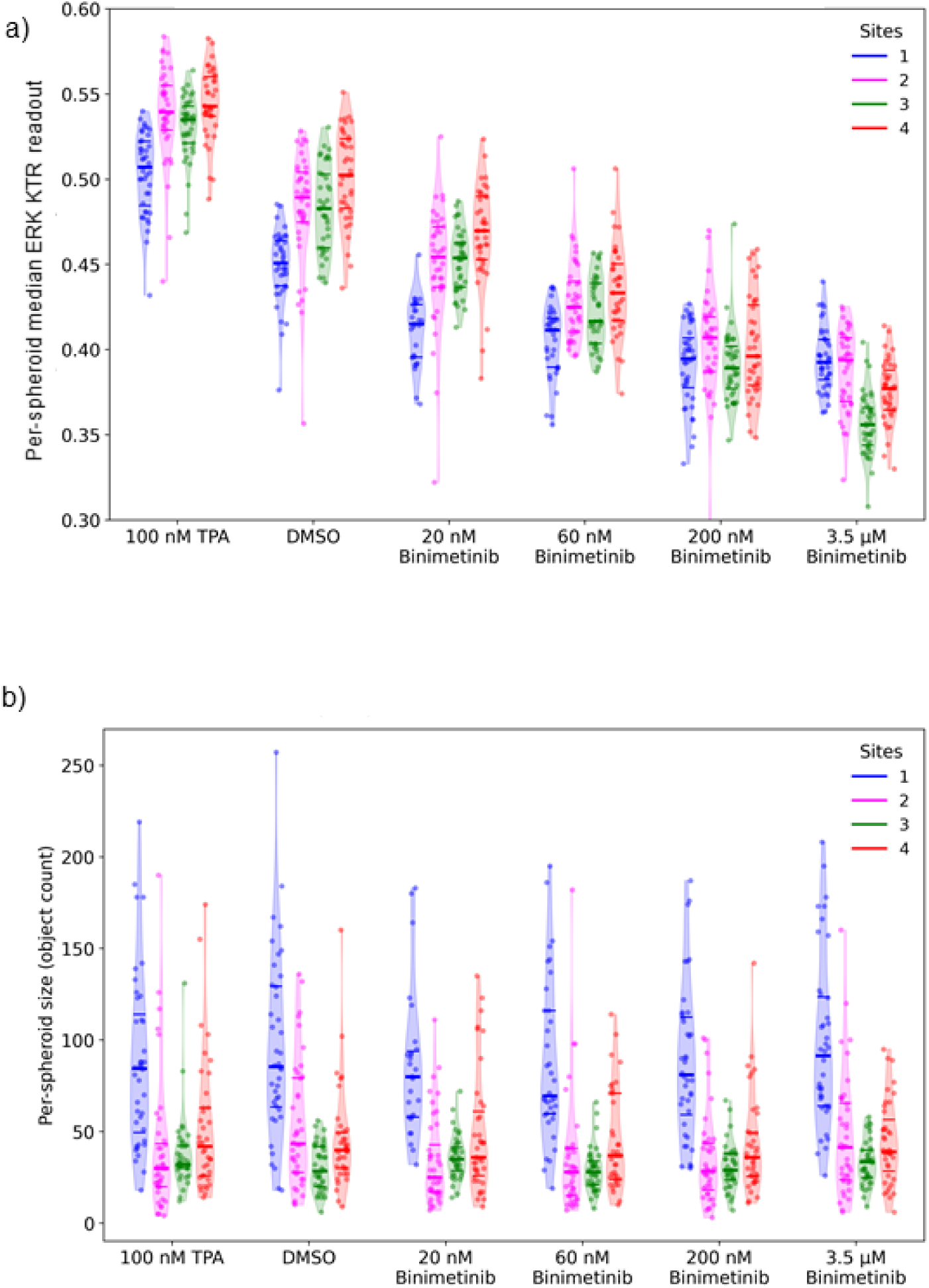
(a) ERK KTR readout and (b) number of cells per spheroid for different conditions and different sites. In (a), points show median of per-cell values in each spheroid and violin plots show the same data as a distribution. b) Points show number of cells per spheroid and violin plots show the same data as a distribution. Thick horizontal bars show median per-spheroid value for each condition/site and thin horizontal bars indicate the inter-quartile range.

We then applied a linear mixed effects model [41] using site 3 and DMSO as reference. Analysis of the 934 spheroid-level observations (target to image and segment 240 spheroids per site, 934 imaged and segmented across all sites) revealed highly significant effects of both condition (ANOVA χ^2^ = 862, degrees of freedom (df) = 5, p < 0.001) and site (χ^2^ = 88, df = 3, p < 0.001), as well as a significant condition-by-site interaction (χ^2^ = 130, df = 15, p < 0.001).

Across all sites, treatment conditions affected the cytoplasm-to-nucleus intensity ratio, with the 3.5 µM Binimetinib condition producing the strongest reduction relative to DMSO control as expected. The overall response profile was broadly consistent across all the sites. Relative to site 3 (intercept M = 0.481, standard error (SE) = 0.005), site 4 showed a statistically significant elevated baseline value (β = 0.017, SE = 0.007, *t*(70.7) = 2.66, *p* = 0.01). Conversely, site 1 exhibited a significantly reduced baseline value (β = −0.042, SE = 0.007, *t*(69.4) = −6.46, p < 0.001). Site 2 did not differ significantly from site 3 (β = −0.004, SE = 0.007, *t*(69.4) = −0.66, *p* = 0.5). While statistically significant, these baseline differences suggest minor variations in fluorophore expression level, fluorescence imaging or segmentation across sites. Furthermore, these differences persisted across all treatment conditions (see below), indicating stable site-specific offsets rather than condition-dependent biological differences

Using the model estimated means we calculated the dynamic range of the assay. For a given site, this was defined as the difference in readout between the maximum (100 nM TPA) and minimum (3.5 µM Binimetinib) biosensor readouts given by equation 1. These results are summarised in the Table 1.

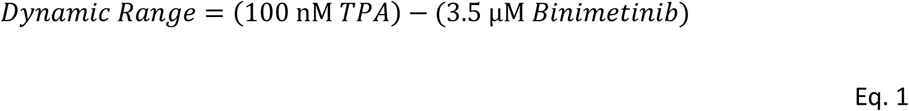

**Table 1.**
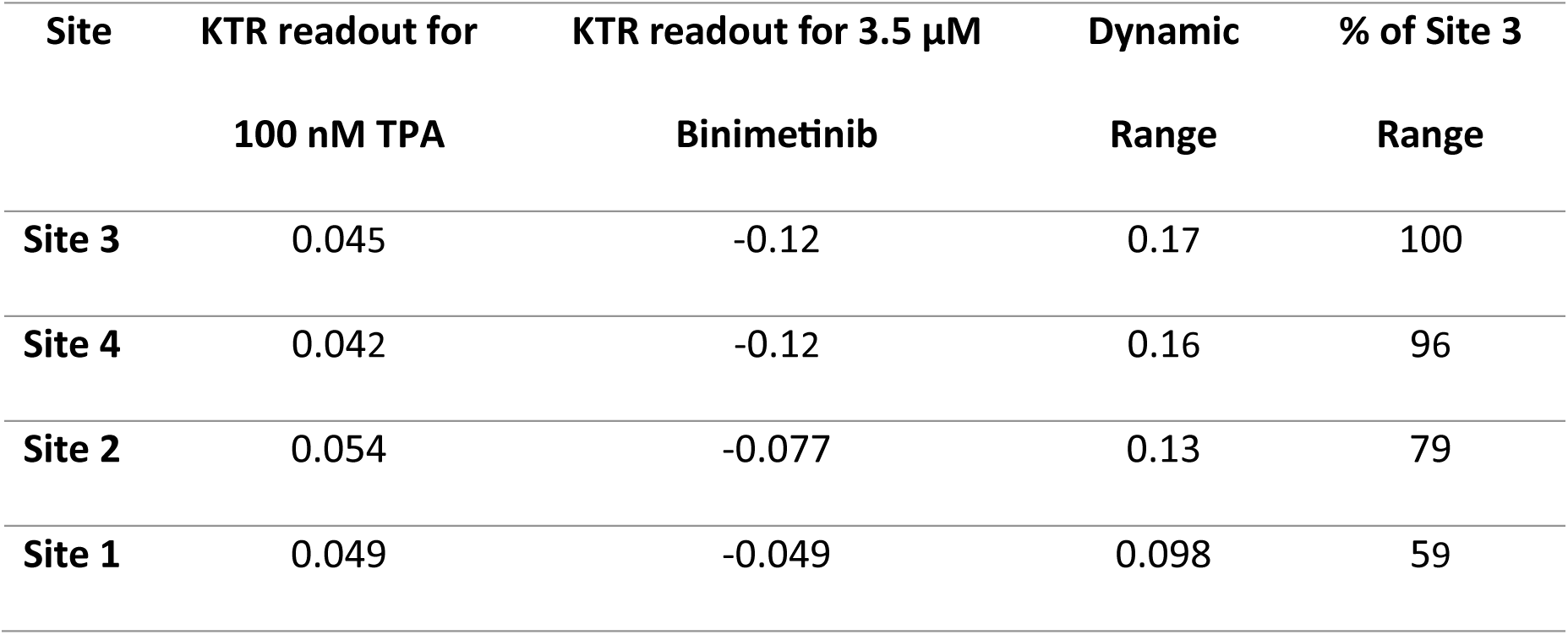
Dynamic range across sites calculated from model estimated means.

Similarly, we also estimated baseline differences expressed as a fraction of the Site 3’s dynamic range, see Table 2.

**Table 2.** Absolute estimated baseline difference expressed as a percentage of reference site range.

| Comparison | Absolute Difference | % of Site 3 Range |
| --- | --- | --- |
| Site 3 vs Site 4 | 0.015 | 9.1 |
| Site 3 vs Site 2 | 0.010 | 6.0 |
| Site 3 vs Site 1 | 0.021 | 12 |

### Condition effects

Binimetinib demonstrated a clear dose-dependent reduction of the cytoplasm-to-nucleus intensity ratio compared to DMSO control. This pattern was consistent across all four sites (all Dunnett’s *p* < 0.001, see Supplementary Table Dunnett’s).

For the 100 nM TPA condition, as expected, the cytoplasm-to-nucleus intensity ratio was elevated relative to the baseline condition (Dunnett’s p < 0.001 for all sites).

### Condition-by-site interactions

While the overall pattern of response to treatments was robust, statistically significant interactions were observed for three conditions. For site 1, statistically significant interactions were observed for the two highest Binimetinib doses (200 nM, β = 0.037, SE = 0.009, *t*(69.4) = 4.01, *p* < 0.001 and 3.5 µM, β = 0.073, SE = 0.009, *t*(69.4) = 7.87, *p* < 0.001), where the effect was smaller than at site 3, suggesting presence of a site-specific effect. For the highest 3.5 µM Binimetinib dose, the interaction between Binimetinib and site 2 was statistically significant (β = 0.044, SE = 0.009, *t*(69.4) = 4.79, *p* < 0.001) with a smaller effect compared to site 3. It is important to note that none of the other condition-by-site interactions were statistically significant.

Finally, the minimal random variance at the well level (SD = 0.0048) indicates low technical noise. The treatment effects were large and consistent across all the sites (all Dunnett *p* <0.001).

## Discussion

In this work, we set out to assess the reproducibility of a 3D spheroid-based assay with sub-cellular KTR readout using dOPM across four separate research sites. Researchers at all sites used the same protocol for preparing the assay and perform the imaging. Below we consider factors that could be controlled better or differently in future experiments, but we are not able to ascertain from the data acquired how much variance can be attributed to each of these factors.

### Reagent differences

Sites purchased reagents from same suppliers with same product codes, so some of the variance between sites may be due to batch-to-batch variability of the reagents.

### Cell culture differences

Our protocol did not define the frequency of cell splitting or trypsinisation duration and temperature used for passaging the cells. Also, the number of cell passages after thawing was not controlled between sites, see Table 6. The higher number of passages at Site 1 may have contributed to the larger spheroids at Site 1.

### Imaging differences

Image acquisition was identical between sites, except for minor variations in the emission filter used when acquiring the H2B-iRFP670 channel that was not noticed until after the experiments were performed. Our protocol of using a single plane of a single spheroid to set the laser power for imaging resulted in a ∼6-fold variation in H2B-iRFP670 signal between the sites with the minimum and maximum raw signals. For the ERK-KTR-mRuby2 channel, this variation was ∼2-fold. In the future, this difference in signal levels between sites could be avoided by sampling a larger range of spheroids when setting the laser powers and this process could be performed automatically for each spectral channel.

### Image analysis differences

The same image analysis algorithm was applied to all datasets, with one person performing all image analysis. Manually determined image processing parameters were established on an initial subset of the data from site 3 and then implemented for the analysis of all data without further adjustment.

While this approach removed inter-operator bias, it is recognised that fixed-parameter pipelines remain sensitive to site-to-site variability. Establishing settings on a specific subset can lead to parameters that are overfitted to the training data, which may not fully generalise across different sites. Future work could address this by using manually annotated training datasets from each site to optimise parameters via grid search, or by adopting foundation models like Cellpose 3 that are pre-trained on large, diverse datasets [42, 43].

## Conclusion

We have studied the reproducibility across four geographically dispersed institutes of a high content spheroid-based assay read out in 3D via dOPM imaging in 96-well plates. Semi-automated imaging and segmentation were applied to analyse the resulting images to quantify a KTR biosensor at the per-cell level. Random variance at the well level was negligible (SD = 0.0048 relative to range of KTR biosensor readout at reference site of 0.17), indicating low technical noise and validating the spheroid-level inference. Treatment effects were dose-dependent and highly significant across all sites compared to DMSO control (Dunnett-corrected *p* < 0.001) despite site-specific modulation of effect magnitudes at extreme doses.

The maximum KTR readout was achieved with treatment by 100 nM TPA and the minimum value by 3.5 μM Binimetinib. We applied a linear mixed-effects model and the range in KTR readout between these two treatments varied between 59% to 96% across sites relative to the reference site (Site 3). For all sites, the differences in KTR readout were statistically significant for all treatments compared to DMSO control value for that site. Site-specific baseline differences were consistent across all treatment conditions, indicating minor variations in fluorophore expression levels, fluorescence imaging, or segmentation. The range in measured bias between sites was between 6 and 12% of the range of the reference site. Site 1 exhibited a stable site-specific statistically significant reduced baseline and site 4 showed a significantly elevated baseline compared to site 3. Overall, our work provides a report on the degree of reproducibility that can be expected from this spheroid-based KTR assay with dOPM readout across independent institutes.

## Methods

### Cell line

The assay was based on the pre-existing mouse NRAS activated melanocyte cell line (19161) [32, 33] modified at the ICR to express an ERK-KTR-mRuby2, EGFP-CAAX and H2B-iRFP670, see Table 3. Fluorescent transgenes were introduced by lentiviral transduction with psPax (packaging) and MD2.G (envelope) plasmids, using Effectene (QIAGEN) transfection reagent.

**Table 3:** Transgenes incorporated into the NRAS cell line.

| Construct | Plasmid description | Source |
| --- | --- | --- |
| ERK-KTR | pLentiPGK Blast DEST<br>ERKKTRmRuby2 | Addgene #90231 |
| EGFP-CAAX | pCSII- IRES2-hygro | Sahai Lab |
| Histone-iRFP670 | pLentiPGK DEST H2B-iRFP670 | <u>Addgene #90237</u> |

### Expansion of cell line and transfer to other sites

The NRAS cell line was expanded at the ICR, frozen in liquid nitrogen and shipped to the other 3 partner sites on dry ice. The cell lines were then thawed and culture established at each site to establish normal proliferation and to check for expression of the FPs.

### Cell culture

Cells were cultured in complete-medium (DMEM with 10% FBS and Penicillin-Streptomycin (10,000 U/mL)). Cells were passaged according to standard practice at each institute. Table 4 lists the cell culture medium components, suppliers and catalogue numbers.

**Table 4:** Cell culture medium components.

| Description | Supplier | Catalogue number |
| --- | --- | --- |
| DMEM-Dulbecco's Modified Eagle Medium | Thermo Fisher Scientific | CAT#41966029 |
| Trypsin-EDTA (0.25%), phenol red | Thermo Fisher Scientific | CAT#25200056 |
| Penicillin-Streptomycin (10,000 U/mL) | Thermo Fisher Scientific | CAT#15140122 |
| Fetal Bovine Serum (FBS), qualified, heat inactivated | Thermo Fisher Scientific | CAT#16140071 |

### Plate preparation

Samples were prepared in multi-well plates according to the plate map shown in Figure 2. A schematic of the final contents of the wells is shown in Figure 1. The reagents used are listed in Table 5.

**Table 5:** Reagents and items for plate preparation. Day 1, plate cells.

| Name | Description | Supplier | Catalogue number |
| --- | --- | --- | --- |
| Multi-well plates | PhenoPlate™, 96-well,<br>black, optically clear<br>flat-bottom, tissue-<br>culture treated, lids,<br>case of 40 | Revvity | 6055302 |
| Cultrex | Reduced growth<br>factor, basement<br>membrane extract<br>Type II, Pathclear | R&D systems | 3533-005-02 |
| MEKi | Binimetinib (MEK162),<br>Synonyms: ARRY-162,<br>ARRY-438162 | Selleckchem | S7007 |
| TPA | Phorbol 12-myristate<br>13-acetate | Sigma | P8189-1mg |
| Fluorescent beads | 200 nm diameter<br>Tetraspek beads | Thermofisher | T7280 |

#### Day 1, plate cells

At ∼70% confluency, cells were harvested via trypsinization (1mL trypsin 0.25%), neutralized with complete medium (5 mL) and centrifuged at 200 × *g* for 5 min. Following resuspension in complete medium (1 mL) cells were counted using the cell counter available at each site.

The required number of cells were centrifuged (200 × *g* for 5 min) and resuspended in complete medium. The BME was kept on ice to avoid premature BME gelation. Working quickly, cells were mixed with BME to achieve a 95% BME suspension with 2 × 10^5^ cells/mL and then dispensed into 24 wells of the 96-well plate at 100 μL/well to achieve 20,000 cells/well. Plates were incubated at 37°C for 35 min to induce gelation, after which 200 μL of 2% BME culture medium was added to each well.

#### Days 2&5, change media

100 μL (half) of existing 2% BME overlying the gelled BME matrix was removed and replaced with 100 μL of fresh 4% BME. Addition of 2x concentration of BME is to compensate for BME consumed and aims to maintain 2% BME in solution overlying gel.

#### Day 8, prepare for imaging

Tetraspeck beads (1 in 20 dilution from stock) in BME gel were prepared to achieve a suspension of beads in 95% BME. 100 μL of bead and BME mixture was pipetted into the bead well with 200 μL 2% BME on top.

For the cell-containing wells, medium was exchanged with 4% BME as described above. Drug treatments were added as 50 μL of 7x desired final concentration and ensuring a constant 0.1% final DMSO concentration across all conditions.

Following the assay, we realised that the protocol used for splitting cells was not uniform across sites. Table 6 provides the details on the splitting procedure at each site and the number of cell passages at the point the assay was run.

**Table 6:** Summary of cell culture conditions and passage numbers used at each site.

| Site | 1 | 2 | 3 | 4 |
| --- | --- | --- | --- | --- |
| Trypsinisation temperature | Room temp. | Room temp. | 37° | 37° |
| Trypsinisation duration (mins) | 2-3 | 2-3 | 5 | 5 |
| Cell splitting frequency (per week) | 2 | 3 | 1 | 1 |
| Cell passages since thaw | 18 | 5 | 9 | 2 |

### Imaging

Prior to imaging, we used a 20x air immersion objective lens (Nikon, CFI Plan Fluor 20× Objective, NA 0.75, MRH00205) and the Nikon Perfect Focus system to automatically generate a map of the z-height of the in-use wells of the 96-well plate.

A 4× magnification objective lens was then used to capture a single brightfield image at the centre of each well to record an overview of the morphology of the spheroids (0.25 GB total data volume).

The pre-find procedure starts with tiled wide-field epifluorescence imaging of the H2B-iRFP670 channel over a 3 × 3 mm² area in spheroid-containing wells using a 20× objective lens with 4×4 on-chip binning, giving an effective pixel size of 1.3 µm. Seven z-planes were acquired at 25 µm intervals in each well starting at the top of the coverslip, generating 2.56 GB of image data across all 24 wells. The resulting z-stacks were processed using custom Python code to identify spheroids suitable for high-resolution dOPM imaging. A fixed camera offset of 1760 DN (110 × 4²) was subtracted to remove the background DC component. Each z-plane was then smoothed separately with a Gaussian filter (σ = 3 px) and a broad uniform filter (window = 80 px). The difference between these two images enhanced locally compact structures while suppressing slowly varying out-of-focus background. A 2-level Otsu threshold was then applied and connected components were labeled and filtered by their volume, retaining multicellular spheroids with effective radii between 20 µm and 200 µm. For each object, the most in-focus plane was defined as the z-slice with the highest median raw intensity sampled through z across the object’s XY region. The in-plane centroid from that slice was converted to stage coordinates using per-plane metadata. Where available, up to ten of the largest spheroids separated from one another by a lateral distance of at least 100 µm were then exported as a Nikon NIS-Elements multipoint position list for automated dOPM acquisition.

dOPM imaging of each spheroid was then performed using a 60× magnification water immersion lens (Nikon, CFI Plan Apo VC 60× water-immersion, NA 1.2, MRD07602) using the system described in [27]. Briefly, the system employs a 50x/0.95 NA secondary/tertiary objective and a tertiary tube lens with a focal length of 50 mm. The pixel size at the sample was 0.35 μm. Two views were acquired of each volume at ±35° and 151 planes were acquired per view with a 1 μm step in the direction perpendicular to the illumination plane (∼100 GB total, for 240 spheroids). Laser powers were adjusted when imaging a plane through a single spheroid (that was not used in subsequent imaging), aiming to achieve >1000 (DN) across the body of the spheroid in each channel. Overall, for each plate we aimed to acquire 240 3D volumes in 3 spectral channels in ∼2 hours.

The dOPM data was then deskewed and fused into the lab frame using bead registration data (acquired separately, see Methods) with a 0.7^3^ μm^3^ voxel size (37 GB total).

Table 7 provides a summary of the image acquisition process and Table 8 summarises the image data acquired.

**Table 7:** Summary of plate imaging process.

| Step | Description | Objective | Details | Approximate duration (minutes) |
| --- | --- | --- | --- | --- |
| 1 | Plate z-map measurement | 20x air | Measure z-position of bottom of 96-well plate using Nikon Perfect Focus System (PFS). Results used to set z position of image stacks acquired in step 3. | 5 |
| 2 | Overview Imaging | 4x air | A single brightfield image at the center of each well to record an overview of the spheroid distribution. | 5 |
| 3 | Widefield pre-find | 20x air | Tile-scan of $3 \times 3 \text{ mm}^2$ area per well in 7 z-planes (25 $\mu\text{m}$ z-step) imaging H2B-iRFP670. Python code then used to detect up to 10 spheroids per well. | 30 |
| 4 | Bead imaging for registration (start) | 60x water | Optimise correction collar, align light-sheet and image bead volumes for view registration. | 10 |
| 5 | dOPM imaging | 60x water | Imaging of 240 spheroids in 3 channels with 2 dOPM views | 120 |
| 6 | Bead imaging for registration (end) | 60x water | Repeat image acquisition from step 4 to verify optical alignment stability. | 5 |

**Table 8:** Summary of image data acquired.

| Parameter | Details |
| --- | --- |
| dOPM views | 2, $\pm 35^\circ$ light-sheet tilt |
| Raw z-planes | 151 (1 $\mu\text{m}$ steps) |
| Raw image size | 512×512 pixels (sensor cropped) |
| Raw pixel size | 0.35×0.35 $\mu\text{m}^2$ (sample space) |
| Resliced voxel size | 0.7×0.7×0.7 $\mu\text{m}^3$ (sample space) |
| Spectral channels | GFP (488 nm ex, 525/45 nm em)<br>KTR (561 nm ex, 609/54 nm em)<br>H2B (642 nm ex, 694/44 nm em, site 1;<br>642 nm ex, 697/58 nm em, sites 2&4;<br>642 nm ex, 698/70 nm em, site 3) |
| Camera exposure time | 10 ms |
| Total acquisition time | 30 seconds per spheroid, ~2 hours overall |
| Total dataset | 10 spheroids per well<br>4 wells per condition<br>6 conditions<br>240 spheroids total (0.454 GB per spheroid, 106 GB total) |

### Image analysis

Image analysis was performed using a variant of the algorithm reported by Ratcliffe et al. [44], which is described below.

For each fluorescence channel, the background was removed by subtracting the mode intensity of each image volume.

Nuclear segmentation was performed using a nonlinear multiscale top-hat enhancement transform following Santos et al.[45] and Guglielmi et al.[46]. This was performed by first smoothing with 3D Gaussian filters at three spatial scales of σ₁ ≈ 0.7 μm, σ₂ ≈ 1.4 μm, σ₃ ≈ 2.8 μm (corresponding to kernel sizes of 1, 2, and 4 voxels at 0.7 μm/voxel), resulting in images U_1_, U_2_ and U_3_ respectively. The enhancement used two paired scale comparisons: the normalized difference between U₁ and U₂, i.e. (U₁−U₂)/U₂, and between U₂ and U₃, i.e. (U₂−U₃)/U₃. These responses were combined linearly with a fixed mixing weight (a₁ = 0.7) to prioritize fine-scale structure. A global threshold of 0.13 was then applied to generate a binary mask.

Small holes were filled and thin structures removed using morphological operations. A spherical morphological opening with a radius of 0.7 μm was used to suppress small bright spots. Objects smaller than ∼2.1 μm in diameter were excluded. A 3D watershed refinement step was applied to separate touching nuclei, using a distance transform of the binary regions followed by 3D Gaussian smoothing (σ = 1.65 μm).

Finally, nuclei were restricted to the main spheroid by dilating and eroding the binary mask with a large spherical structuring element (∼15 μm radius) to merge nearby regions, identifying the largest connected component, and retaining only nuclear labels overlapping this component.

For quantifying the KTR readout, a cytoplasmic “collar” region was defined for each nucleus by performing two successive 1 μm 3D dilations of the nuclear mask. The collar was then extracted as the shell-like region between the second dilation and the original nucleus, excluding any areas overlapping nuclear or collar regions from neighbouring cells. The mean KTR intensities were then quantified separately within nuclear and cytoplasmic compartments and the final KTR readout is calculated as the mean cytoplasmic to mean cytoplasmic plus mean nuclear ratio.

### Statistical analysis

To assess the reproducibility of our 3D fluorescence microscopy pipeline across four independent sites, we fitted linear mixed-effects models to spheroid-level cytoplasm-to-nucleus ratio values. Mixed effects models were chosen to explicitly account for the hierarchical structure of the data, wherein, spheroids were nested within wells and wells were nested within sites. As cells within a spheroid share the same microenvironment and are likely to be clonal, the inference was performed on spheroid level. For each spheroid, the mean cytoplasm-to-nucleus ratio (mean_cnr) was computed across all segmented cells and used as the dependent variable.

The full model included fixed effects for condition and site, with their interaction term to evaluate whether drug responses varied systematically across sites. Random intercepts were specified for wells nested within sites to account for technical replicates. All models were fitted using the lme4 package in R (R-Studio 4.5.0) [47], with p-values for fixed effects obtained using Satterwhaite’s approximation as implemented in lmerTest [48]. The model specification was as follows:

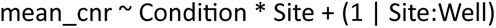

We used maximum likelihood estimation (REML = FALSE) for this model to enable direct assessments of fixed effects. This approach provides estimates of variance components while accounting for the hierarchical structure of the experimental design. Type III analysis of variance (ANOVA) was used to test main effects and interactions. Post-hoc pairwise comparisons of each treatment condition to the DMSO control (within each site) were conducted using Dunnett’s method, which is optimal for control-group designs and controls family-wise error rate. All tests used α = 0.05; estimated degrees of freedom were computed using Satterthwaite’s approximation (fixed effects) and Kenward-Roger’s approximation (ANOVA).

## Data Availability

Datasets reported in this manuscript are available from the BioImage Archive under accession S-BIAD3591: https://www.ebi.ac.uk/biostudies/bioimages/studies/S-BIAD3591. The software used for image processing and analysis is available as permanent archived snapshots on Zenodo: Fiji plugin scripts for dOPM data processing https://doi.org/10.5281/zenodo.20849636, software steps used in the multisite assay https://doi.org/10.5281/zenodo.20932316. The live development version of the analysis code is available on GitHub: https://github.com/ImperialCollegeLondon/oblique-plane-microscopy/tree/main/dOPM-MultiSiteAssayPaper.

## Acknowledgments

The work reported here was undertaken as part of the MACH3Cancer Accelerator project led by Paul French. The authors gratefully acknowledge the expert technical assistance of Martin Kehoe and Simon Johnson of the Optomechanical Workshop, Department of Physics, Imperial College London. The authors also acknowledge support from the Light Microscopy Facility at the Institute of Cancer Research managed by Kai Betteridge.

## Supplementary figure captions

Supplementary Figure 1. Montage of orthogonal maximum intensity projections (MIPs) of spheroids from a control well from each site for the H2B and KTR channels. Scale bar 100 μm.

Supplementary Figure 2: Graphical representation of spheroid size and ERK KTR readout. All cells from all spheroids in a given condition/site were projected onto the XY plane using spheroid-centered coordinates. The XY space was divided into a 45×45 grid of spatial bins; bins containing fewer than 3 cells were excluded. ERK activity is expressed as the delta versus a site-matched DMSO baseline, with the median ERK KTR readout across DMSO-condition cells computed per site and subtracted from each cell’s raw ratio prior to binning. The diverging colour scale is symmetric around zero with limits set to the 98th percentile of absolute delta values in both directions; values exceeding this range are displayed at the colour limit (saturated).

Supplementary Figure 3. (a) Median H2B-iRFP670 nuclear intensity, (b) median KTR-mRuyby2 nuclear intensity and (c) median KTR-mRuby2 cytoplasmic collar intensity over all cells per condition. Error bars show 95% confidence intervals.

Supplementary Table 1

Pairwise contrasts comparing each treatment condition to DMSO control within each site. Results from a linear mixed-effects model (mean_cnr ∼ Condition × Site + (1 | Site:well)) with Dunnett’s method for multiple comparisons correction (α = 0.05).

Key: Estimate, (β coefficient, difference in KTR readout from DMSO); SE, standard error; df, degrees of freedom (estimated by Kenward-Roger method); Lower CL, Upper CL, 95% confidence limits (Dunnett-adjusted); t ratio, t-statistic (estimate); p value, Dunnett-adjusted p-value.

Supplementary Table 2

Pairwise contrasts between all sites (with site 3 as the reference). Results from the linear mixed effects model with Tukey’s method for multiple comparisons correction (α = 0.05).

Key: Estimate (β coefficient, difference in baseline KTR readout between sites), SE (standard error), df (degrees of freedom, estimated by Kenward-Roger method), Lower CL and Upper CL (95% confidence limits, Tukey-adjusted), t ratio (t-statistic), p value (Tukey-adjusted p-value).

