## Supplementary information for "Multi-Site Reproducibility Study of 3D High-Content Analysis with Dual-View Oblique Plane Microscopy"

### Supplementary Figure 1

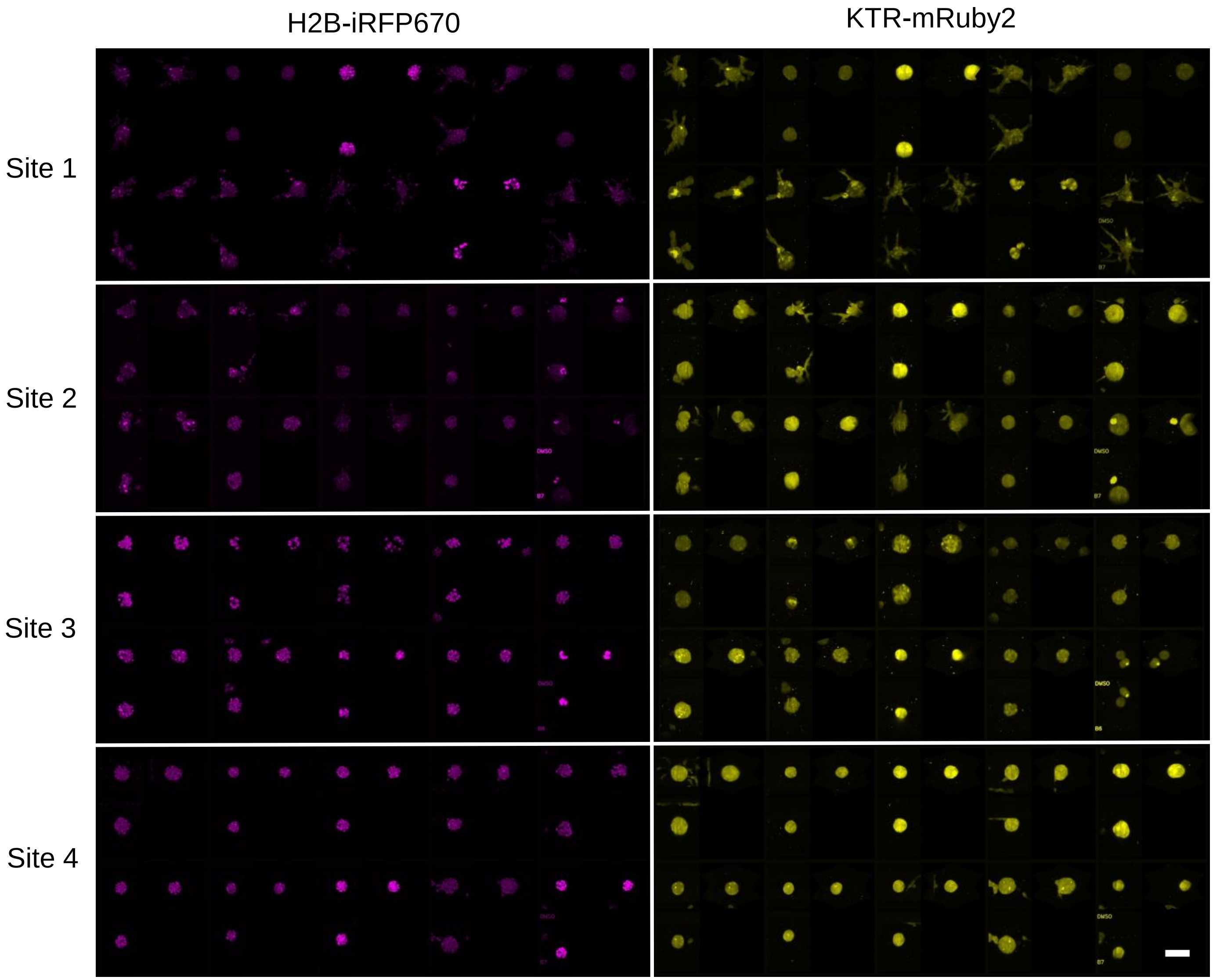

### Supplementary Figure 2

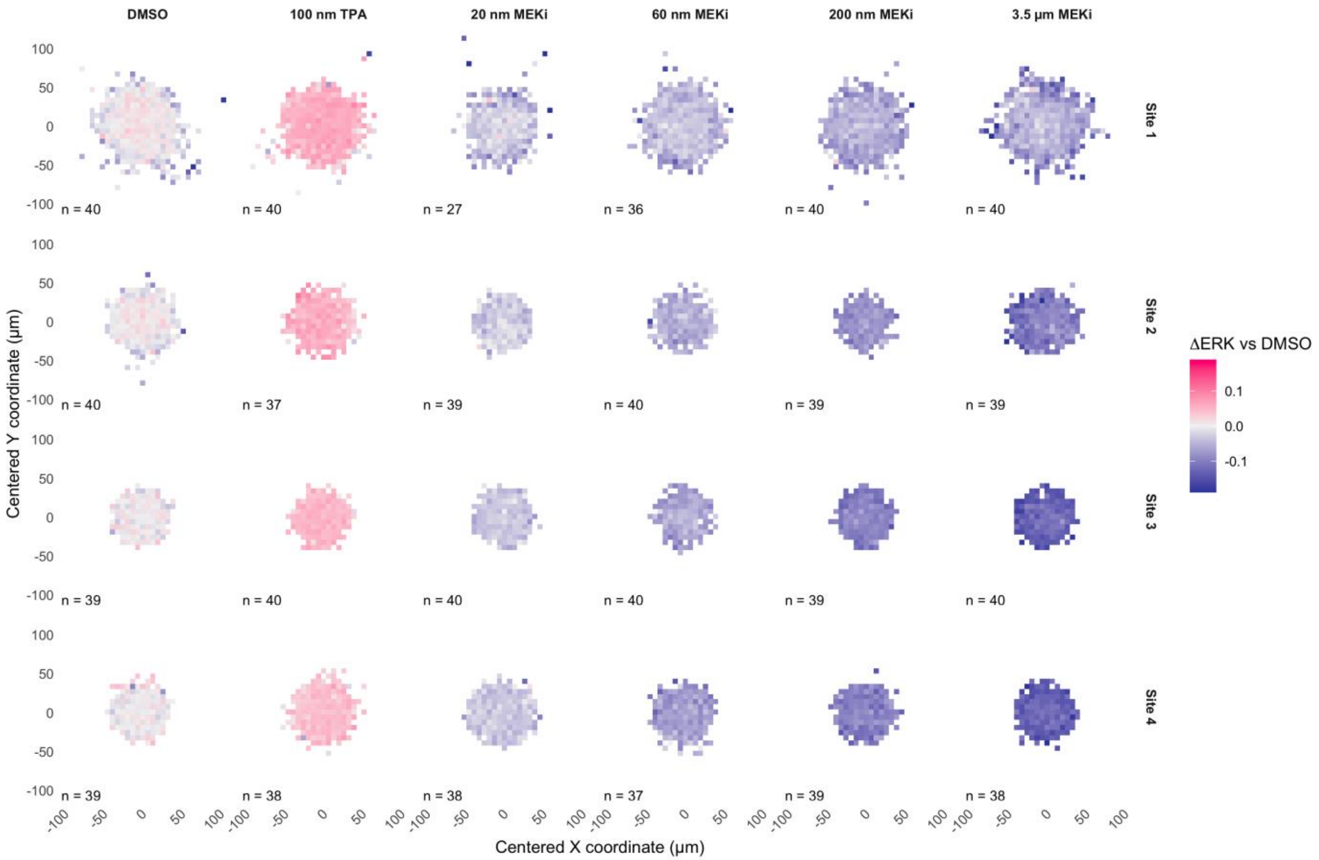

### Supplementary Figure 3

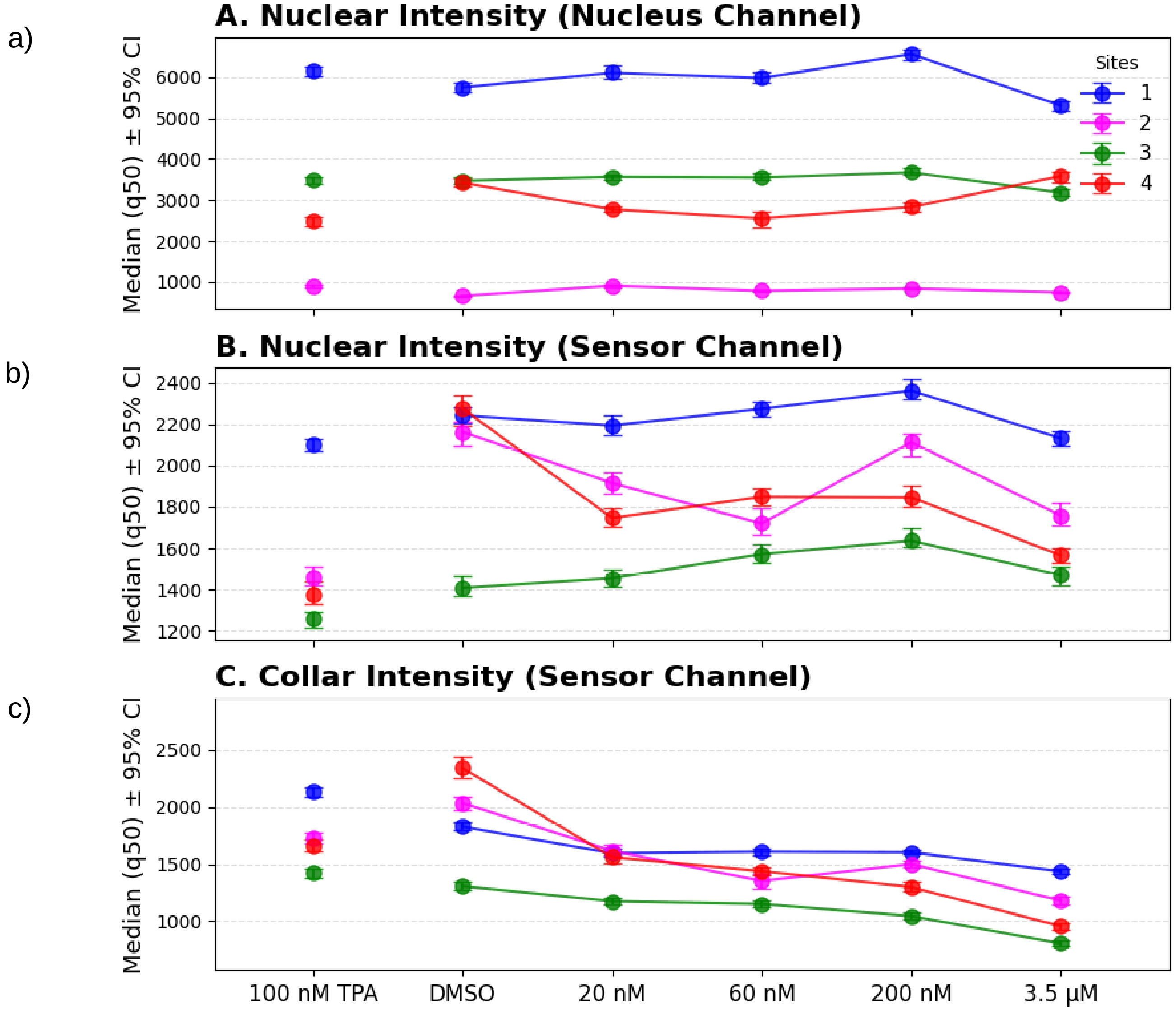

### Supplementary Table 1 – Dunnett comparisons

| contrast | Site | estimate | SE | df | Lower CL | Upper CL | T ratio | p value |
|---|---|---|---|---|---|---|---|---|
| 20 nM Binimetinib - DMSO | 3 | -0.0305 | 0.006512 | 69.11387 | -0.04735 | -0.01365 | -4.68325 | 6.69E-05 |
| 60 nM Binimetinib - DMSO | 3 | -0.05859 | 0.006512 | 69.11387 | -0.07544 | -0.04175 | -8.99724 | 2.98E-13 |
| 200 nM Binimetinib - DMSO | 3 | -0.08922 | 0.006512 | 69.11387 | -0.10606 | -0.07237 | -13.6993 | 0 |
| (100 nM TPA +ve control) - DMSO | 3 | 0.0449 | 0.006512 | 69.11387 | 0.028052 | 0.061748 | 6.894487 | 1.02E-08 |
| (3.5 µM Binimetinib -ve control) - DMSO | 3 | -0.12151 | 0.006512 | 69.11387 | -0.13836 | -0.10466 | -18.6584 | 0 |
| 20 nM Binimetinib - DMSO | 4 | -0.03271 | 0.006575 | 71.62624 | -0.0497 | -0.01571 | -4.97489 | 2.13E-05 |
| 60 nM Binimetinib - DMSO | 4 | -0.06552 | 0.00664 | 74.49267 | -0.08267 | -0.04837 | -9.86818 | 3.55E-15 |
| 200 nM Binimetinib - DMSO | 4 | -0.09426 | 0.006575 | 71.62624 | -0.11126 | -0.07727 | -14.3376 | 0 |
| (100 nM TPA +ve control) - DMSO | 4 | 0.04165 | 0.006607 | 72.99751 | 0.024581 | 0.058719 | 6.304268 | 9.81E-08 |
| (3.5 µM Binimetinib -ve control) - DMSO | 4 | -0.11762 | 0.006575 | 71.62624 | -0.13462 | -0.10063 | -17.8907 | 0 |
| 20 nM Binimetinib - DMSO | 2 | -0.02797 | 0.006512 | 69.11387 | -0.04482 | -0.01113 | -4.29538 | 0.000271 |
| 60 nM Binimetinib - DMSO | 2 | -0.04585 | 0.006512 | 69.11387 | -0.0627 | -0.02901 | -7.04112 | 5.51E-09 |
| 200 nM Binimetinib - DMSO | 2 | -0.07257 | 0.006512 | 69.11387 | -0.08942 | -0.05572 | -11.143 | 0 |
| (100 nM TPA +ve control) - DMSO | 2 | 0.054164 | 0.006512 | 69.11387 | 0.037316 | 0.071012 | 8.316961 | 5.2E-12 |
| (3.5 µM Binimetinib -ve control) - DMSO | 2 | -0.07741 | 0.006512 | 69.11387 | -0.09426 | -0.06056 | -11.8863 | 0 |
| 20 nM Binimetinib - DMSO | 1 | -0.03364 | 0.007125 | 88.10625 | -0.05197 | -0.0153 | -4.72112 | 4.32E-05 |
| 60 nM Binimetinib - DMSO | 1 | -0.04034 | 0.00665 | 74.28232 | -0.05751 | -0.02316 | -6.0654 | 2.52E-07 |
| 200 nM Binimetinib - DMSO | 1 | -0.05231 | 0.006512 | 69.11387 | -0.06915 | -0.03546 | -8.03184 | 4.71E-11 |
| (100 nM TPA +ve control) - DMSO | 1 | 0.049075 | 0.006512 | 69.11387 | 0.032227 | 0.065923 | 7.535523 | 6.59E-10 |
| (3.5 µM Binimetinib -ve control) - DMSO | 1 | -0.04901 | 0.006512 | 69.11387 | -0.06586 | -0.03216 | -7.52566 | 6.88E-10 |

### Supplementary Table 2 – Tukey site contrasts

| Contrast between sites | estimate | SE | df | Lower CL | Upper CL | t ratio | p value |
|---|---|---|---|---|---|---|---|
| 3 - 4 | -0.01513 | 0.002678 | 71.06299 | -0.02218 | -0.00809 | -5.64996 | 1.84E-06 |
| 3 - 2 | -0.00995 | 0.002659 | 69.11387 | -0.01695 | -0.00295 | -3.74256 | 0.002069 |
| 3 - 1 | 0.020606 | 0.002711 | 73.13913 | 0.013478 | 0.027734 | 7.600223 | 3.76E-10 |
| 4 - 2 | 0.005181 | 0.002678 | 71.06299 | -0.00187 | 0.012226 | 1.934473 | 0.223091 |
| 4 - 1 | 0.035737 | 0.00273 | 75.11559 | 0.028563 | 0.042911 | 13.08926 | 0 |
| 2 - 1 | 0.030557 | 0.002711 | 73.13913 | 0.023428 | 0.037685 | 11.2702 | 0 |
